# A survivin-driven tumour-activatable minicircle system for prostate cancer theranostics

**DOI:** 10.1101/2020.06.25.171645

**Authors:** TianDuo Wang, Yuanxin Chen, David Goodale, Alison L. Allan, John A. Ronald

**Affiliations:** Department of Medical Biophysics, Schulich School of Medicine & Dentistry, Western University, London, ON, Canada; Department of Anatomy & Cell Biology, Schulich School of Medicine & Dentistry, Western University, London, ON, Canada; Department of Oncology, Schulich School of Medicine & Dentistry, Western University, London, ON, Canada; Robarts Research Institute – Imaging Laboratories, London, ON, Canada; London Regional Cancer Program, London Health Science Centre, London, ON, Canada; Lawson Health Research Institute, London, ON, Canada

**Keywords:** minicircle, survivin, urinary reporter, prodrug therapy, prostate cancer

## Abstract

Gene vectors driven by tumour-specific promoters to express reporter genes and therapeutic genes are an emerging approach for improved cancer diagnosis and treatment. Minicircles (MCs) are shortened plasmids stripped of prokaryotic sequences and have potency and safety characteristics that are beneficial for clinical translation. We previously developed survivin-driven, tumour-activatable MCs for cancer detection via a secreted blood reporter assay. Here we present a novel theranostic system for prostate cancer featuring a pair of survivin-driven MCs, combining selective detection of aggressive tumours via a urinary reporter test and subsequent tumour treatment with gene-directed enzyme prodrug therapy.

**Methods:** We engineered both diagnostic and therapeutic survivin-driven MCs expressing Gaussia luciferase, a secreted reporter that is detectable in the urine, and cytosine deaminase:uracil phosphoribosyltransferase fusion, respectively. Diagnostic MCs were evaluated in mice carrying orthotopic prostate tumours with varying survivin levels, measuring reporter activity in serial urine samples. Therapeutic MCs were evaluated in mice receiving prodrug using bioluminescence imaging to assess cancer cell viability over time.

**Results:** Diagnostic MCs revealed mice with aggressive prostate tumours exhibited significantly higher urine reporter activity than mice with non-aggressive tumours and tumour-free mice. Combined with 5-fluorocytosine prodrug treatment, therapeutic MCs resulted in reduced bioluminescence signal in mice with aggressive prostate tumours compared to control mice.

**Conclusion:** Sequential use of these MCs may be used to first identify patients carrying aggressive prostate cancer by a urinary reporter test, followed by stringent treatment in stratified individuals identified to have high-risk lesions. This work serves to highlight tumour-activatable MCs as a viable platform for development of gene-based tumour-activatable theranostics.

## Introduction

Gene transfer, the introduction of foreign genetic material into patient cells, is a promising avenue for both novel cancer therapeutics and diagnostics. A shared goal of this field is to develop technologies which promote high on-tumour and minimal-to-no off-tumour transgene expression. One way to achieve this is to create gene vectors driven by tumour-specific promoters to selectively activate transgene expression in cancer cells, but not healthy cells [1]. Historically, these “tumour-activatable” vectors have predominantly encoded therapeutic transgenes for cancer gene therapy [2-5]. However, more recently, researchers have also created tumour-activatable vectors encoding secreted and/or imaging reporter genes as a means for specifically detecting and/or locating cancer cells throughout the body [6-8]. Naturally, gene vectors encoding both reporter and therapeutic transgenes also offer strategies to provide treatment post-diagnosis, extending gene transfer technologies into the exciting field of theranostics [9].

Amongst the numerous discovered tumour-specific promoters, the survivin promoter (pSurvivin) is a promising candidate for use in tumour-activatable vectors. Also known as baculoviral inhibitor of apoptosis repeat-containing 5 (BIRC5), survivin has been widely studied as a cancer biomarker in adults due to its near exclusive presence in cancers cells, but not healthy cells [10-12]. These characteristics have led many groups to leverage the tumour-specific activity of pSurvivin to drive tumour-specific transgene expression in prostate [13], breast [14], and liver cancer [15], amongst others. Furthermore, survivin is elevated in many aggressive, metastatic tumours compared to non-aggressive, indolent tumours [16, 17], a feature that could lead to higher transgene expression in tumours that are the most relevant to patient outcome.

Despite continued refinement, arguably the goal of developing a highly efficient and safe tumour-activatable vector has yet to be fully realized. The majority of tumour-activatable vectors developed thus far have used viral vectors such as lentiviruses and adenoviruses, or non-viral, plasmid constructs. Viral vectors have been the most popular due to their relatively high gene transfer efficiency compared to plasmids [18]. However, random genomic integration by some viruses can disrupt key genes, which raises major concerns regarding patient safety [19]. Additionally, pre-existing immunity and/or unwanted immune responses triggered by viral vectors have often undermined these strategies during clinical trials [20, 21]. Despite continued refinement of viral vectors to be non-integrating and less immunogenic [22], many have posited plasmids may be more favorable for widespread clinical utility in terms of safety [23]. However, the main challenges that recombinant plasmids face are inefficient delivery and transfection, fast clearance, and host incompatibility of the prokaryotic sequences such as selection markers and replication elements [24]. In particular, transfer of antibiotic resistance genes to host microbiota is a biosafety hurdle [25]. The goal of addressing the limitations of plasmids motivated the inception of minicircle (MC) vectors, which are smaller plasmid derivatives stripped of their prokaryotic backbone [26]. Removal of the backbone confers greater transfection efficiency with MCs compared to plasmids through reduction in vector size. Additionally, MCs exhibit prolonged expression profiles compared to plasmids, as the plasmid backbone is a common target for transcriptional silencing [27, 28]. Thus, tumour-activatable MCs could overcome the limitations of tumour-activatable plasmids by achieving improved transfer efficiency into cancer cells, inducing robust and longer tumour-specific transgene expression, while also eliminating prokaryotic components that could otherwise compromise clinical safety and translation into humans.

Our group previously developed the first tumour-activatable MC system that used pSurvivin to drive expression of a secreted reporter gene for cancer detection [29]. These MCs encoded a secreted blood-detectable reporter called secreted embryonic alkaline phosphatase (SEAP). SEAP is a variant of placental alkaline phosphatase that is only expressed during embryogenesis and thus exhibits minimal background activity in antenatal humans [30]. MCs were complexed with a linear polyethylenimine (PEI) transfection agent for intravenous delivery and we were able to reliably distinguish mice with melanoma lung tumours from tumour-free mice through a SEAP blood test [29]. Recently, we have also demonstrated the utility of these MCs for discerning between human prostate cancer xenografts of varying aggressiveness, with higher SEAP blood levels being measured in mice carrying aggressive versus non-aggressive primary prostate tumours [31].

Due to the inherent modularity of this system, here we explore the use of tumour-activatable MCs for cancer theranostics. Specifically, we built and validated a pair of novel survivin-driven MCs, which we have termed diagnostic MCs and therapeutic MCs, that can be used separately or together for cancer detection with a urinary reporter test and/or cancer treatment using gene-directed enzyme prodrug therapy (GDEPT) [32]. We first engineered survivin-driven diagnostic MCs encoding Gaussia luciferase (GLuc), a reporter gene that is isolated from the marine copepod *Gaussia princeps* [33]. Unlike SEAP which is not detectable in the urine, nearly 90% of GLuc that is secreted is cleared via the renal pathway [34]. These GLuc-expressing MCs present several advantages over our original design. Firstly, a urinary-based test better facilitates longitudinal study over blood-based tests due to the ease of urine collection. Secondly, the GLuc gene (∼550 bp) is considerably shorter than SEAP (∼1500 bp), further reducing overall MC size. Moreover, non-human-derived enzyme-based reporter genes, such as GLuc, offer the distinct advantage of being naturally absent from the human body and provide amplified readouts compared to measuring of endogenous compounds [4] – traits which could improve the detection of smaller tumours for earlier diagnoses [35]. For therapeutic MCs, we engineered survivin-driven MCs encoding the fusion enzyme cytosine deaminase:uracil phosphoribosyltransferase (CD:UPRT) [36]. CD:UPRT catalyzes the conversion of the non-toxic prodrug 5-fluorocytosine (5-FC) to the anti-tumour metabolites 5-fluorouracil (5-FU) and 5-fluorouracil monophosphate (5-FUMP), leading to inhibition of cancer proliferation through thymidine deprivation [37]. Importantly, CD:UPRT has been previously used in preclinical models for treating several types of cancer including prostate cancer [38]. Here we demonstrate the complementarity of our diagnostic MCs for detection of aggressive, high-survivin orthotopic prostate tumours in mice via increased urine GLuc activity, along with therapeutic MCs encoding CD:UPRT for attenuating the growth of high-survivin, GLuc-detectable prostate tumours.

## Methods

### Parental Plasmid and MC Construction

A dual-transgene tumour-activatable theranostic parental plasmid (PP) driven by pSurvivin and encoding both GLuc2 and CD:UPRT separated by the P2A self-cleavage peptide sequence (pSurvivin-GLuc2-P2A-CD:UPRT-PP) was designed in-house and built by Genscript (NJ, USA). Single-transgene pSurvivin-driven PPs were designed and made in-house using In-Fusion HD Cloning Kits (Takara Bio, CA, USA). First, pSurvivin-SEAP-PP [29] was digested with AgeI and NheI, and the linearized product without the SEAP transgene was isolated via gel extraction. To create pSurvivin-GLuc-PP, the GLuc2 transgene from pCMV-GLuc2 (NEB, MA, USA) was subcloned into the linearized product. To make pSurvivin-CD:UPRT-PP, the CD:UPRT gene from pSELECT-CD:UPRT (Invivogen, CA, USA) was subcloned into the same linearized product. MCs were generated from their respective PP using a previously described production system [39]. Briefly, each PP was transformed into ZYCY10P3S2T E. coli and viable Kanamycin-resistant colonies were selected and cultured at 37°C in lysogeny broth (LB) for 6 h followed by terrific broth (TB) for 12 h. To generate MCs via site-specific recombination, induction of ϕC31 integrase and Sce-I endonuclease was achieved through addition of equal volume LB containing 0.001% (v/v) L-arabinose and 4 mL 1N NaOH, and incubated for 5.5 h at 30°C. Production of PPs followed the same protocol without the addition of arabinose. PPs and MCs were purified from E. coli using an endotoxin-free Maxi kit (Qiagen, ON, Canada) and resuspended in nuclease-free water. MCs were further cleaned to remove any PP contamination using Plasmid-Safe ATP Dependent DNase kit (Epicenter, WI, USA) followed by the DNA Clean & Concentrator Kit (Zymo Research, CA, USA).

### Cell Culture and Transduction

Human LNCaP PCa cell lines were obtained from ATCC (VA, USA) and PC3MLN4 cells were a kind gift from Dr. Hon Leong (Western University, ON, Canada). Primary prostate epithelial cells were also obtained from ATCC. LNCaP and PC3MLN4 cells were maintained in RPMI 1640 medium (Wisent Bioproducts, QC, Canada). Primary cells were grown in Prostate Epithelial Cell Basal Medium (ATCC). All media was supplemented with 10% (v/v) Fetal Bovine Serum (FBS) and 5% (v/v) Antibiotic-Antimycotic and cells were cultured at 37°C in 5% CO_2_. The absence of mycoplasma contamination was routinely verified using the MycoAlert Mycoplasma Detection Kit (Lonza, NY, USA). To generate cells that can be detected using bioluminescence imaging, PC3MLN4 and LNCaP naïve cells were transduced with lentiviral vectors (in 8 μg/mL polybrene) encoding tdTomato (tdT) and Firefly luciferase (FLuc) driven by the human elongation factor-1 alpha promoter (pEF1α). Transduced cells were washed, and tdT-positive cells were sorted using a FACSAria III fluorescence-activated cell sorter (BD Biosciences, CA, USA).

### In Vitro Assessment of Diagnostic MCs

PCa and primary prostate epithelial cells were seeded at 5×10^4^ cells/well in a 24-well plate one day prior to transfection. Cells were transfected with pSurvivin-GLuc-MCs (1 μg) after complexation with 2 μL of jetPEI, a linear polyethylenimine transfection agent (Polyplus Transfection, PA, USA). On day 2 post-transfection, 100 μL of media was collected from each well, centrifuged at 10,000 x g for 10 min, and supernatant was stored at -20°C. Each time media was collected, wells were washed with Phosphate Buffered Saline (PBS) and fresh media was added; thus, GLuc measurement reflected protein production over the desired time intervals. GLuc activity in samples was measured using the Biolux Gaussia Luciferase Assay kit (NEB, Ipswich, MA). GLuc assay solution (50 μL) was added to 20 μL of each sample and total luminescence over 5s (expressed in relative light units, RLU) was measured using a Glomax 20/20 luminometer (Promega, WI, USA). Separately seeded cells were transfected with pCMV-GLuc2 plasmids (1 μg, NEB) and GLuc activity was measured using the same kit. GLuc activity from pSurvivin-GLuc-MCs was normalized to GLuc activity from pCMV-GLuc plasmids. To compare expression from MCs against their PP counterparts, PC3MLN4 cells were seeded at 5×10^4^ cells/well in a 24-well plate. One day later, cells were either transfected with equimolar pSurvivin-GLuc-PPs (1 μg) or pSurvivin-GLuc-MCs (0.42 μg) complexed with jetPEI. To maintain the same mass of DNA and volume of jetPEI between wells, pEF1α-tdt-luc2-PPs (0.58 μg) were co-transfected into cells receiving MCs. GLuc activity in media samples were assessed as described above. To visualize intracellular GLuc activity, media was removed from transfected cells, washed with D-PBS, and fresh media was added prior to addition of 5μΜ h-coelenterazine to each well. Plates were then imaged on an IVIS Lumina XRMS scanner (PerkinElmer, MA, USA). Average radiance (p/s/cm^2^/sr) per well was quantified by placing regions-on-interest over each well using Living Image 4.5.2 software (PerkinElmer, MA, USA).

### In Vitro Assessment of Therapeutic MCs

To evaluate therapeutic MCs, PCa cells were seeded in 24-well plates and transfected with pSurvivin-CD:UPRT-MCs (1 μg). Cells were transferred to 6-well plates one day later, cell media was supplemented with 500 μg/mL 5-FC (Sigma, MO, USA). For control conditions, either pSurvivin-CD:UPRT-MCs and/or 5-FC in media was withheld. Every other day, media in wells were removed and replaced with fresh media with 500 μg/mL 5-FC. We evaluated the percentage of dead cells using the Zombie Violet Fixable Viability Kit (Biolegend, CA, USA). Briefly, on day 5 post-transfection, cells were collected and resuspended in PBS containing 1:500 Zombie Violet dye. Following 15 min incubation in the dark, the percentage of dead cells (dye positive cells at 405 nm) was determined using a FACSCanto flow cytometer (BD Biosciences). Flow cytometry results were analyzed using FlowJo v10. To measure viability of FLuc^+^ cells using BLI, the same steps of seeding, transfection, and passaging steps were used as above. FLuc BLI was done on days 0, 2, 4, 5, 6, and 7 on the IVIS scanner after addition of 150 μg/mL D-luciferin to each well and average radiance (p/s/cm^2^/sr) for each well was measured. After each imaging session, wells were washed with PBS and fresh media containing 500 μg/mL 5-FC was added.

### In Vitro Assessment of Theranostic MCs

To assess GLuc activity and cell death from PC3MLN4 cells transfected with dual-transgene pSurvivin-GLuc2-P2A-CD:UPRT-MCs, the same procedures were used as described for diagnostic and therapeutic MCs, respectively.

### Tumour Models

All animal procedures were approved by the University Council on Animal Care at the University of Western Ontario (Protocol #2015-058) and are in compliance with Canadian Council on Animal Care (CCAC) and Ontario Ministry of Agricultural, Food and Rural Affairs (OMAFRA) guidelines. For all animal work, 6-8 weeks old male nu/nu athymic nude mice were used (Charles River Laboratories, QC, Canada). For subcutaneous tumours models (n=5), one million PC3MLN4 cells were injected subcutaneously into the right flank of mice. Tumour volume was assessed using calipers and MCs were injected intratumorally once tumours reached ∼150mm^3^. Tumour volume was calculated using the formula [40]: *Tumour volume = 0*.*5*(length x width*^*2*^). For orthotopic tumours models (n=18), mice were anesthetized, maintained at 2% isoflurane, and a small incision (< 1 cm) was made along the midline to expose the lower peritoneum. Gently lifting the bladder from the abdominal cavity towards the head, the prostate was located, and one million PC3MLN4 FLuc^+^ cells were injected into the right anterior prostate lobe. An identical procedure was performed using LNCaP FLuc^+^ cells mixed with equal volume Matrigel (VWR, ON, Canada) to aid tumour formation [41]. Incisions were closed with sutures and surgical staples, and mice were administered Metacam analgesic. BLI was performed weekly post-surgery on the IVIS scanner to assess of FLuc^+^ tumour development and tumours were injected intratumorally with MCs ∼3-6 weeks post-surgery. As a rough estimate for tumour size, orthotopic PC3MLN4 tumour (n=9) volumes were measured using calipers prior to MC-injections, and tumour volume was correlated to BLI signal which was measured one day prior to surgery. At endpoint, animals were euthanized using an overdose of isofluorane followed by cervical dislocation. Tumour tissue was dissected and cut into two parts. One part was immersed in 4% paraformaldehyde (PFA) for 24 h at 4°C then immersed in PBS at 4°C and the second part was snap-frozen with liquid nitrogen then stored at -80°C.

### In Vivo Diagnostic MC Assessment in Subcutaneous and Orthotopic Tumours

Diagnostic MCs were prepared by combining pSurvivin-GLuc2-MCs (25 μg/mouse) with 3 μL of in vivo-jetPEI (Polyplus Transfection) to achieve an N/P ratio of 8 (N/P ratio describes the number of nitrogen residues in in vivo-jetPEI per nucleic acid phosphate). This MC-PEI complex was then resuspended in equal volumes of 10% (w/v) glucose and incubated at room temperature for 30 min. For subcutaneous PC3MLN4 tumours (n=5), mice were anesthetized and maintained with 2% isoflurane, MC complexes were injected intratumourally into 3-4 loci. For intramuscular injections in healthy mice (n=4), MC complexes were injected directly into the right flank. For these animals, urine was collected one day pre-and on days 2, 5, and 7 post-MC injection. Urine collection was done by placing individual mice into small, sterile boxes and each mouse was monitored every 5 min for urination. Urine samples (∼50-100 μL/mouse) were pipetted into a microcentrifuge tube, centrifuged briefly, and then cleared supernatant was stored at -20°C until assayed. Urine GLuc activity was assessed using the same kit as with cell culture media. For orthotopic tumours, animals were anesthetized and maintained with 2% isoflurane, a small incision (< 1 cm) was made along the midline to expose the lower peritoneum, and the orthotopic tumour was located. MC complexes were then injected into 3-4 loci of each intraprostatic tumour (PC3MLN4, n=10; LNCaP, n=3), then incisions were closed with sutures and staples. Tumour-free mice received equivalent injections of MC complexes into the anterior prostate lobe (n=6). Urine samples was collected one day pre- and daily post-MC injection for 7 days. To evaluate the cumulative secretion of GLuc into urine over the 7-day period, area-under-curve (AUC) analysis of GLuc activity measurements taken over time was performed.

### In Vivo Therapeutic MC Assessment in Orthotopic Tumours

Therapeutic MCs were prepared by combining pSurvivin-CD:UPRT-MCs (50 μg/mouse) with 6 μL vivo-jetPEI to achieve an N/P ratio of 8 and injected intratumourally as described above for diagnostic MCs. Mice with PC3MLN4 FLuc^+^ orthotopic tumours were administered with either MC-PEI complexes (n=9) or an equal volume of 0.9% saline as a control (n=9). All animals received intraperitoneal injections of 500 mg/kg 5-FC diluted in 0.9% saline daily from days 1-7 post-injection, then every other day from days 7-14 post-injection. BLI was performed on days 0, 7 and 14 post MC-injection to assess cancer cell viability. BLI images were obtained using an automatic exposure time (max. 60s) until peak signal was reached. Total tumour flux (p/s) at peak signal was quantified by manually placing regions-on-interest over the primary tumour.

### Western Blot Analysis

Snap-frozen tumours were digested with radioimmunoprecipitation assay lysis buffer containing phosphatase inhibitors (Sigma). Protein concentrations were measured with a Pierce BCA protein assay (23227, Thermo Scientific, MA, USA) and 50 μg of protein per sample was separated on 12% SDS-PAGE then electro-blotted onto nitrocellulose membranes using the iBlot Dry Blotting System (Thermo Scientific). Membranes were blocked with 3% bovine serum albumin in PBS with 0.02% Tween 20 (PBST) for 1 h and then probed with rabbit anti-survivin antibody (1:10000 dilution; Ab 76424, Abcam) at 4°C overnight (16 h). Following three washes with PBST, the membrane was incubated with IRDye 800CW goat anti-rabbit IgG (1:5000 dilution, P/N 925-32211, LI-COR, NE, USA) for 1 h. Expression of GAPDH was used as a loading control using a mouse anti-GAPDH antibody (1:10000 dilution, MAB374, Sigma) and IRDye 680CW goat anti-mouse IgG secondary antibody (1:5000 dilution, P/N 926-32220, LI-COR). Fluorescent signal was measured using the Odyssey Imaging System (LI-COR). The same protocol was used to measure survivin levels in cell lysates.

### Statistical Analysis

All statistical analyses were performed in GraphPad Prism 8.0 software. Data were expressed as mean ± SD. For *in vitro* and *in vivo* studies, a Student’s t-test was used to measure differences between two groups and a one-way ANOVA followed by Tukey’s multiple comparisons test was used to compare means for more than two groups. When comparing multiple group means over time *in vivo*, a repeated measures two-way ANOVA followed by Sidak’s multiple comparisons test was used. For all tests, a nominal p-value less than 0.05 was deemed significant.

## Results

### In vitro evaluation of all-in-one MCs co-expressing reporter and therapeutic genes

As the first step towards establishing a tumour-activatable theranostic MC system, we co-encoded both GLuc and CD:UPRT transgenes on a single pSurvivin-driven construct to create an “all-in-one” PP (pSurvivin-GLuc2-P2A-CD:UPRT-PPs; 8.1 kb; Fig. 1A) and subsequent “all-in-one” theranostic MCs (4.1 kb). Proper MC production was confirmed by gel electrophoresis (Fig. 1B). All-in-one MCs were complexed with PEI and transfected into PC3MLN4 cells, which we have previously shown to express a high level of survivin [31]. Prior to transfection, no appreciable levels of GLuc activity was found in media, but significantly greater GLuc activity above baseline was detected two days post-transfection (Fig. 1C). To evaluate the therapeutic effect, the percentage of dead cells was determined in transfected or naïve cells with or without 5-FC treatment using flow cytometry of Zombie violet stained cells. This revealed 27.4 ± 4.5% of cells were dead 4 days after cells were transfected with all-in-one MCs and treated with 5-FC, which was significantly higher compared to all other conditions.

**Figure 1.**
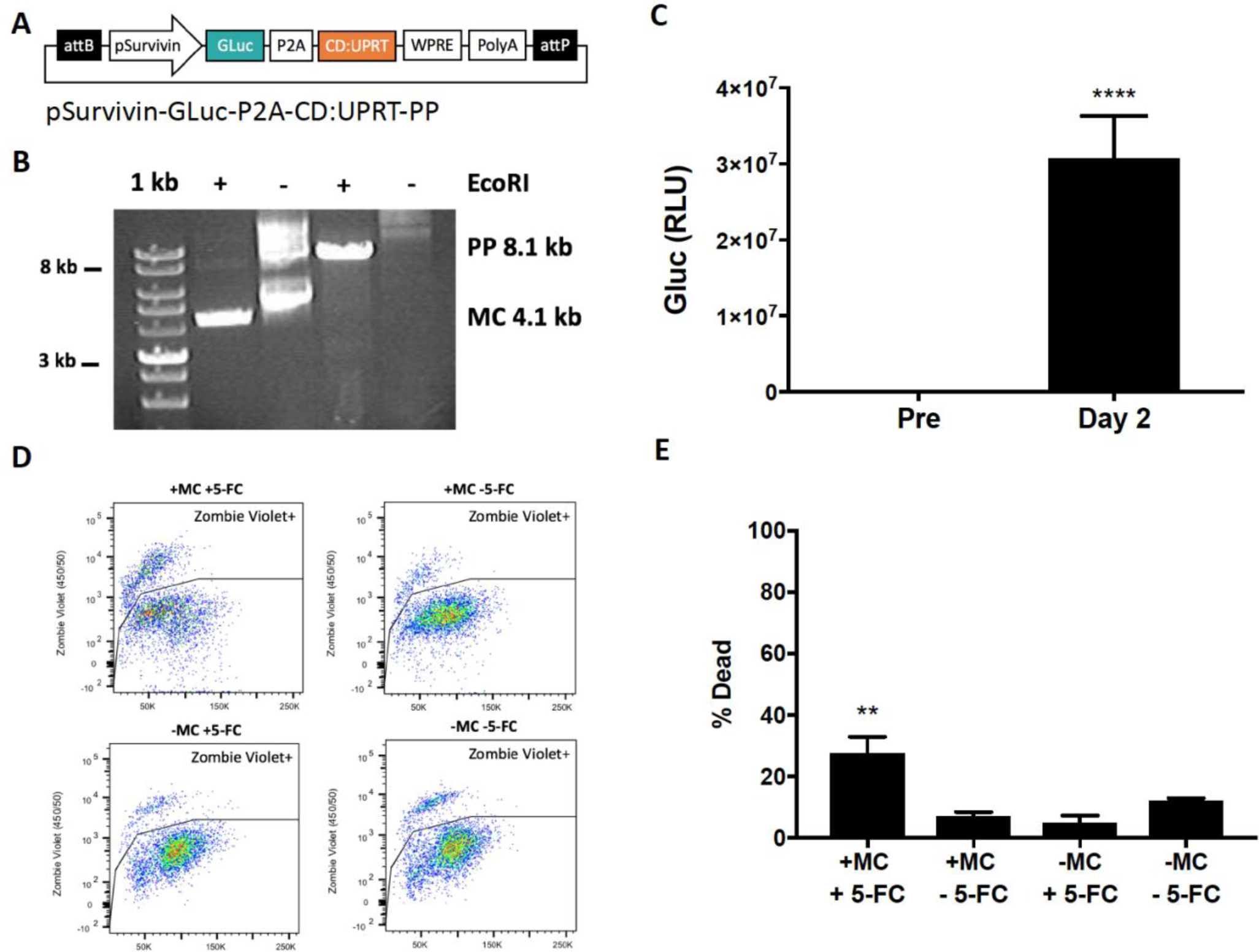
Characterization of theranostic MCs in vitro. (A) Vector map of pSurv-GLuc-CD:UPRT-PPs expression cassette with (B) agarose gel electrophoresis to confirm proper production of PP (8.1 kb) and MC (4.1 kb). (C) GLuc activity in cell supernatant on pre and post-transfection with pSurv-GLuc-CD:UPRT-MCs (n=5). (D) Flow cytometry plots of PC3MLN4 cells on day 5 post-transfection stained with the Zombie Violet Cell Fixable Viability Kit with (E) quantification of Zombie Violet+ (dead) cells (n=3). Data are presented as mean ± SD (\*\*\*\**p* < 0.001, \*\**p* < 0.01).

### Diagnostic MCs induce Gaussia luciferase activity related to cellular survivin levels

Although all-in-one MCs showed potential as a theranostic in survivin-rich PCa cells, we posited that a dual MC system composed of two smaller MCs encoding GLuc and CD:UPRT separately would improve transfection efficiency and transgene expression, potentially leading to increased diagnostic and therapeutic efficacy. We first engineered diagnostic PPs (pSurvivin-GLuc-PP; 6.9 kb) and successfully produced diagnostic MCs (2.9 kb; Fig. 2A and B). Equimolar amounts of MCs and their PP counterparts were transfected into PC3MLN4 cells and after two days, MCs produced significantly higher GLuc activity in culture media than PPs (Fig. S1). Compared to all-in-one MCs (Fig. 1B), diagnostic MCs displayed increased GLuc activity at similar timepoints. We also used BLI to visualize intracellular GLuc activity from diagnostic MCs across PCa cells with varying survivin levels (Fig. 2C). Two days post-transfection, PC3MLN4 cells exhibited significantly higher (>100-fold) signal above background signal than LNCaP cells (∼10-fold; Fig. 2D, indicative of significantly greater GLuc expression in PC3MLN4 (survivin-high) cells compared to LNCaP (survivin-low) cells. To evaluate GLuc secretion, two days post-transfection, GLuc activity in culture media was significantly higher in PC3MLN4 cells, compared to both LNCaP and primary prostate epithelial cells (Fig. 2E). No significant difference in GLuc activity between LNCaP and primary prostate cells was found. To account for variable transfection efficiency across cell types, cells were also transfected with constitutively-on pCMV-GLuc plasmids. Comparable GLuc activity between PC3MLN4 and LNCaP cells was found, while primary prostate cells exhibited significantly lower (∼60%) GLuc activity (Fig. 2F). GLuc activity values obtained using the pCMV-GLuc plasmids were used to normalize GLuc values obtained with the pSurvivin-GLuc-MCs (Fig. 2G). Following normalization, PC3MLN4 cells maintained significantly higher GLuc activity values compared to both LNCaP and primary prostate cells, reflective of the relative survivin levels across the cell types.

**Figure 2.**
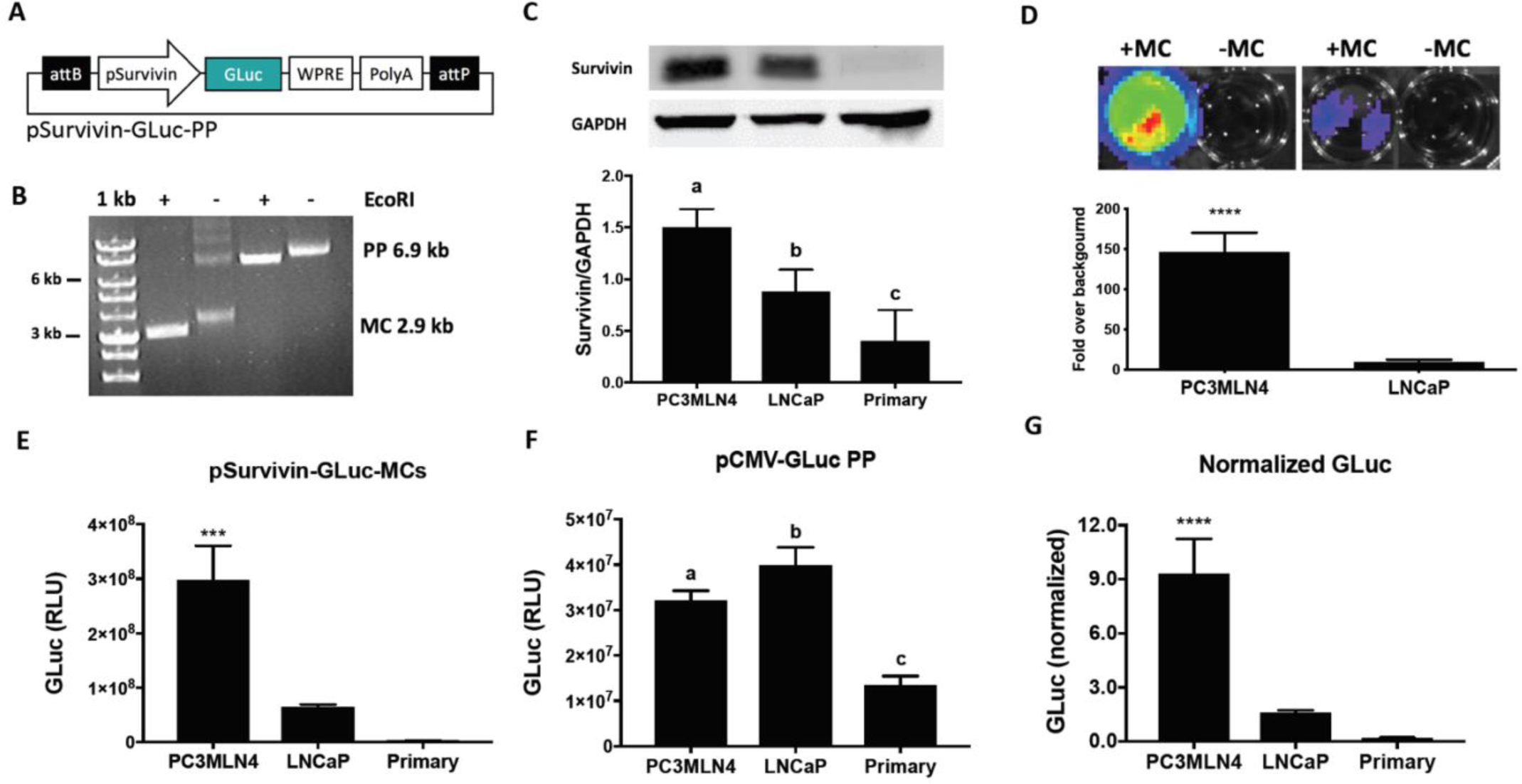
Characterization of secreted reporter expression in vitro. (A) Vector map of pSurv-GLuc-PPs expression cassette with (B) agarose gel electrophoresis to confirm proper production of PP (6.9 kb) and MC (2.9 kb). (C) Western blot for cellular survivin expression in PCa and primary prostate epithelial cells (n=3). Band intensities were quantified using ImageJ and shown relative to GAPDH. (D) GLuc bioluminescence signal above background from PCa cells transfected with pSurv-GLuc-MCs (n=5). GLuc activity in supernatant from cell lines transfected with either (E) pSurv-GLuc-MCs or (F) pCMV-GLuc plasmids on day 2 post-transfection with (G) normalized values (n=3). (H) Groups designated by different letters are significantly different (*p* < 0.05). Data are presented as mean ± SD (\*\*\*\**p* < 0.001, \*\*\**p* < 0.005).

### Diagnostic MCs identify mice carrying aggressive prostate tumours via increased urine GLuc activity

To first evaluate diagnostic MCs *in vivo*, we established PC3MLN4 FLuc^+^ subcutaneous tumours on the right flank of male nude mice and performed intratumoural injections with pSurvivin-GLuc-MCs (25 μg). Urine was collected one day before and then on days 2, 5, and 7 post-MC injection. In tumour-bearing mice, urine GLuc activity was significantly increased on all days post-MC injection (peaking on day 2) compared to GLuc activity pre-MC as well as at all time points following intramuscular MC injections (Fig. S2).

To assess diagnostic MCs across different tumour types, we established orthotopic PC3MLN4 FLuc^+^ and LNCaP FLuc^+^ prostate tumours. A linear correlation was found between *in vitro* FLuc BLI signal and viable cell number in these engineered cells (Fig. S3). As the same number of LNCaP cells displayed roughly 6-fold lower BLI signal than PC3MLN4 cells, we adjusted the signal expected for LNCaP tumours accordingly when tracking tumour growth. Once tumours were ∼150mm^3^ (3-4 weeks for PC3MLN4, 5-6 weeks for LNCaP, Fig. S4), diagnostic MCs (25 μg) were injected intratumourally and urine was collected both one day prior and daily post-MC administration (Fig. 3A). Prior to MC injection, all animals exhibited negligible GLuc activity in urine. For mice carrying PC3MLN4 tumours, urine GLuc activity peaked on day 2, which was significantly higher compared to mice with LNCaP tumours and tumour-free mice at this same timepoint (Fig. 3C). After day 2, GLuc activity in PC3MLN4 mice declined by ∼60% compared to peak day 2 levels and remained at a steady low level until endpoint (Fig. 3D). In mice carrying LNCaP tumours and tumour-free mice which received intraprostatic MC injections, urine GLuc activity post-MC injection was not significantly above baseline levels throughout the study. Area-under-curve (AUC) analysis of GLuc activity over time showed significantly higher AUC values in mice carrying PC3MLN4 tumours compared to all other groups (Fig. 3E). Endpoint Western blot analysis of tumour tissue revealed significantly increased survivin levels in PC3MLN4 compared to LNCaP tumours (Fig. 3F).

**Figure 3.**
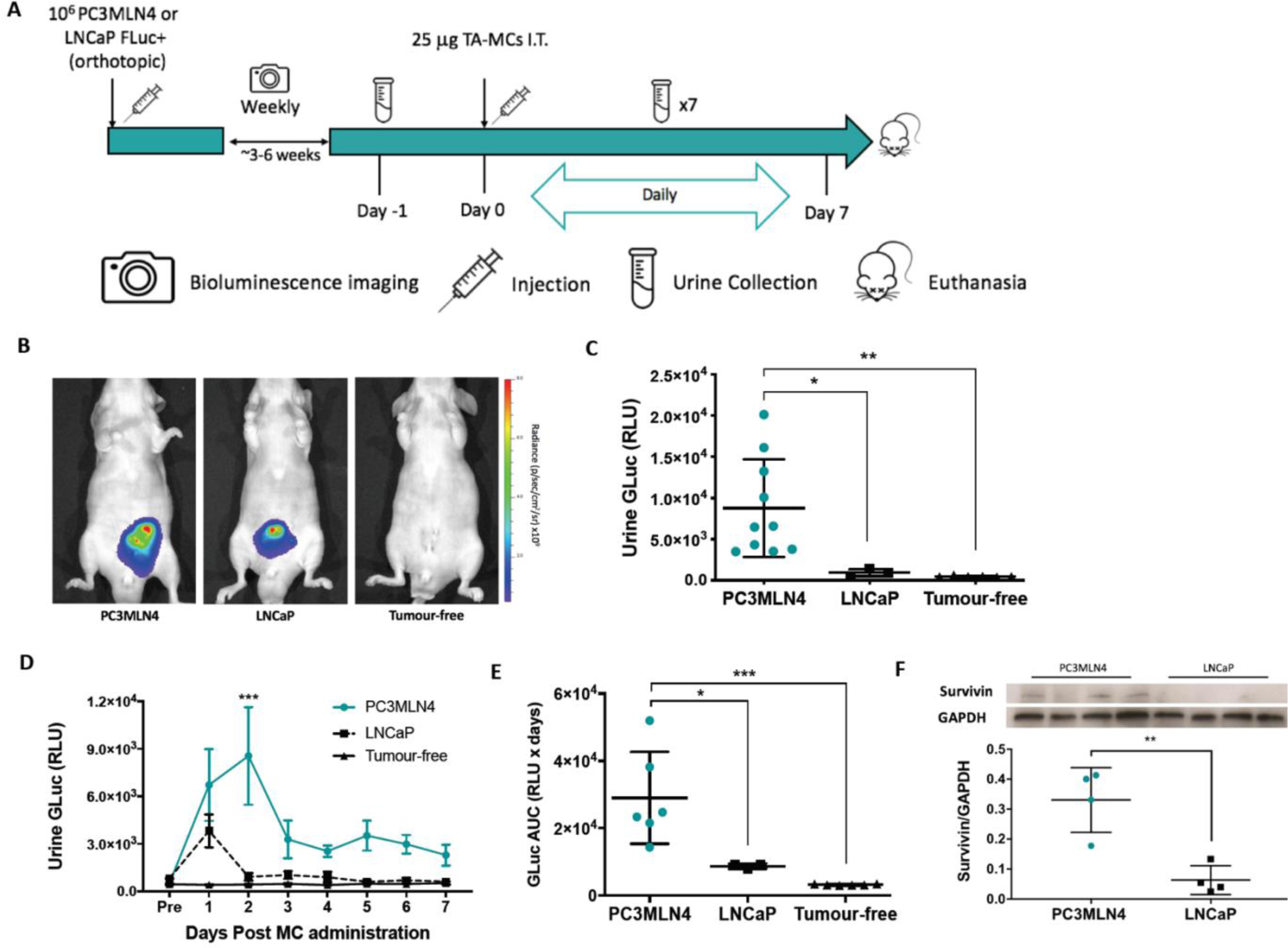
Evaluation of urine GLuc activity in nude mice with orthotopic prostate cancer tumours. (A) Timeline of intratumoural administration of pSurv-GLuc MCs and subsequent urine sampling. (B) Representative FLuc BLI images of mice 30 days after tumour implantation. (C) Urine GLuc activity on day 2 post-MC administration PC3MLN4 (n=10), LNCaP (n=3) or no tumours (n=6) (D) Daily urine GLuc activity from mice bearing PC3MLN4 (n=6), LNCaP (n=3) or no tumours (n=6), and (E) AUC analysis over one week post-MC administration from mice bearing PC3MLN4 (n=6), LNCaP (n=3) or no tumours (n=6). (F) Western blot for survivin levels in tumour lysates (n=4). Data are presented as mean ± SD (\*\*\**p* < 0.005, \*\**p* < 0.01, \**p* < 0.05).

### Therapeutic MCs are selectively cytotoxic in survivin-high prostate cancer cells

To evaluate the use of pSurvivin-driven MCs for therapy, we engineered pSurvivin-CD:UPRT-PPs (7.5 kb) and successfully produced therapeutic MCs (3.5 kb, Fig. 4A and B). Cell death post-transfection with pSurvivin-CD:UPRT-MCs was assessed using flow cytometry of Zombie violet stained cells and 46.9 ± 6.9% of PC3MNL4 cells transfected and treated with 5-FC were dead, and this percentage was significantly higher than all other conditions (Fig. 4C and E). In contrast, no differences in the percentage of dead cells were found for LNCaP cells between all treatment groups (Fig. 4D and F). These findings suggest that therapeutic MCs are selectively cytotoxic to PCa cells expressing high survivin levels when supplied with the prodrug 5-FC under the conditions used here, although we hypothesize cytotoxicity can be optimized by improving transfection efficiency and increase prodrug dosage. Based on these results, we furthered explored the effects of therapeutic MCs on PC3MLN4 FLuc^+^ cells by performing BLI to evaluate cell viability. By day 4 post-transfection and beyond, PC3MLN4 FLuc^+^ cells treated with 5-FC exhibited significantly reduced BLI signal compared to all other conditions (Fig. S5A and B).

**Figure 4.**
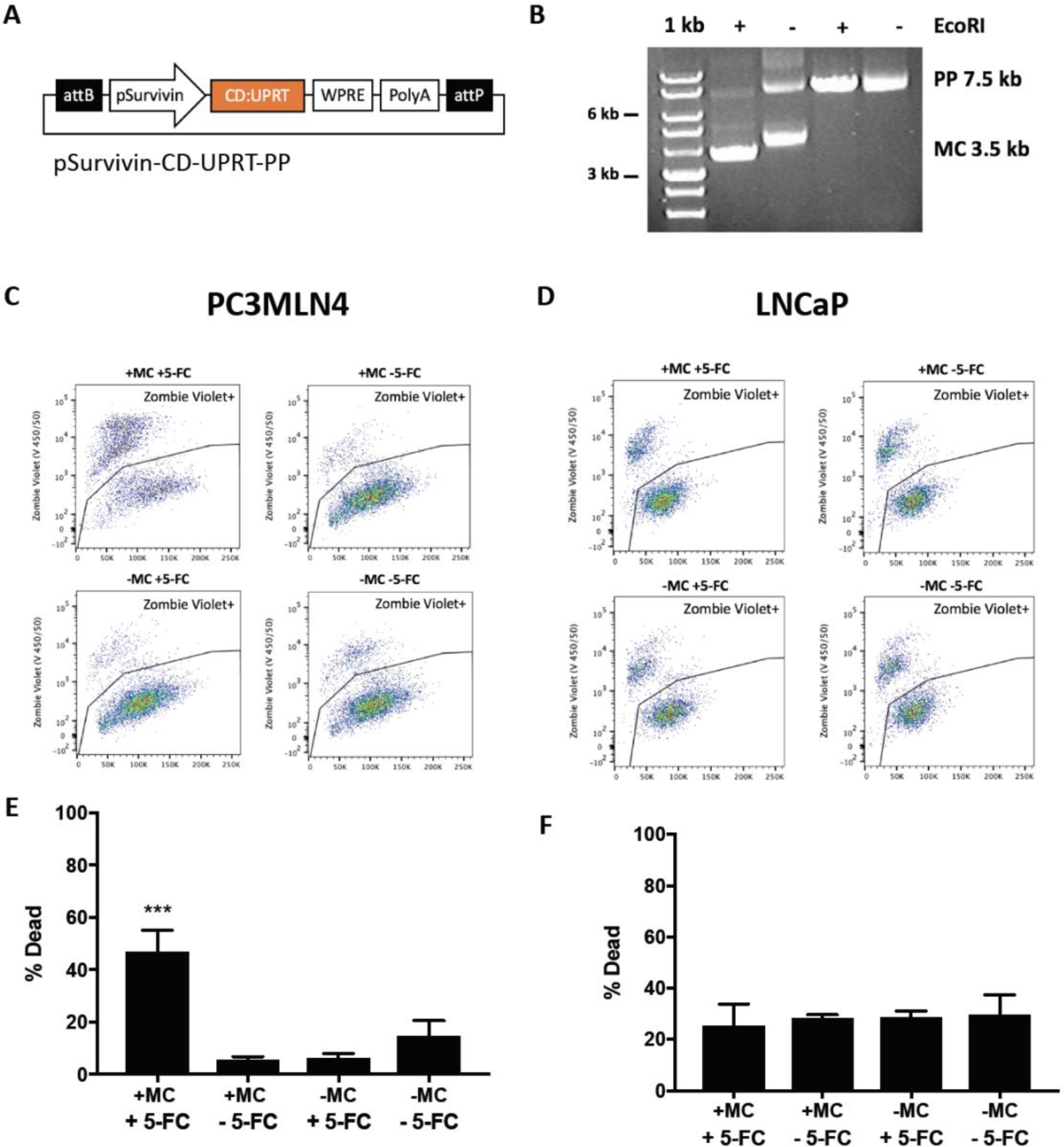
Characterization of suicide gene therapy system on PCa cells with varying survivin. (A) Vector map of pSurv-CD:UPRT-PPs expression cassette with (B) agarose gel electrophoresis to confirm proper production of PP (7.5 kb) and MC (3.5 kb). Flow cytometry plot of (C) PC3MLN4 and (D) LNCaP cells on day 5 post-transfection stained with the Zombie Violet Cell Fixable Viability Kit with (E,F) quantification of Zombie Violet+ (dead) cells (n=3). Data are presented as mean ± SD (\*\*\**p* < 0.005).

### Therapeutic MCs attenuated the growth of aggressive orthotopic prostate cancer

We next evaluated the effects of therapeutic MC administration on the growth of survivin-rich orthotopic PC3MLN4 FLuc^+^ tumours in nude mice. To mitigate discrepancies in initial tumour size between groups, tumour burden was assessed weekly with BLI and once BLI signal reached ∼10^11^ p/s, therapeutic MCs (50 μg) or saline was injected intratumorally. On the day of MC injection (day 0), tumour BLI signal was not significantly different between groups (Fig. 5B and C). Following MC or saline injections, mice were treated daily with 500 mg/kg 5-FC over 14 days (Fig. 5A) and cancer cell viability was monitored over time with BLI. On average, saline-treated mice showed a significant 3.16-fold increase in BLI signal over the 14-day treatment period, while BLI signal in mice receiving therapeutic MCs did not significantly change (Fig. 5B and C). At endpoint, mice which received therapeutic MCs exhibited significantly lower tumour BLI signal than saline-treated mice.

**Figure 5.**
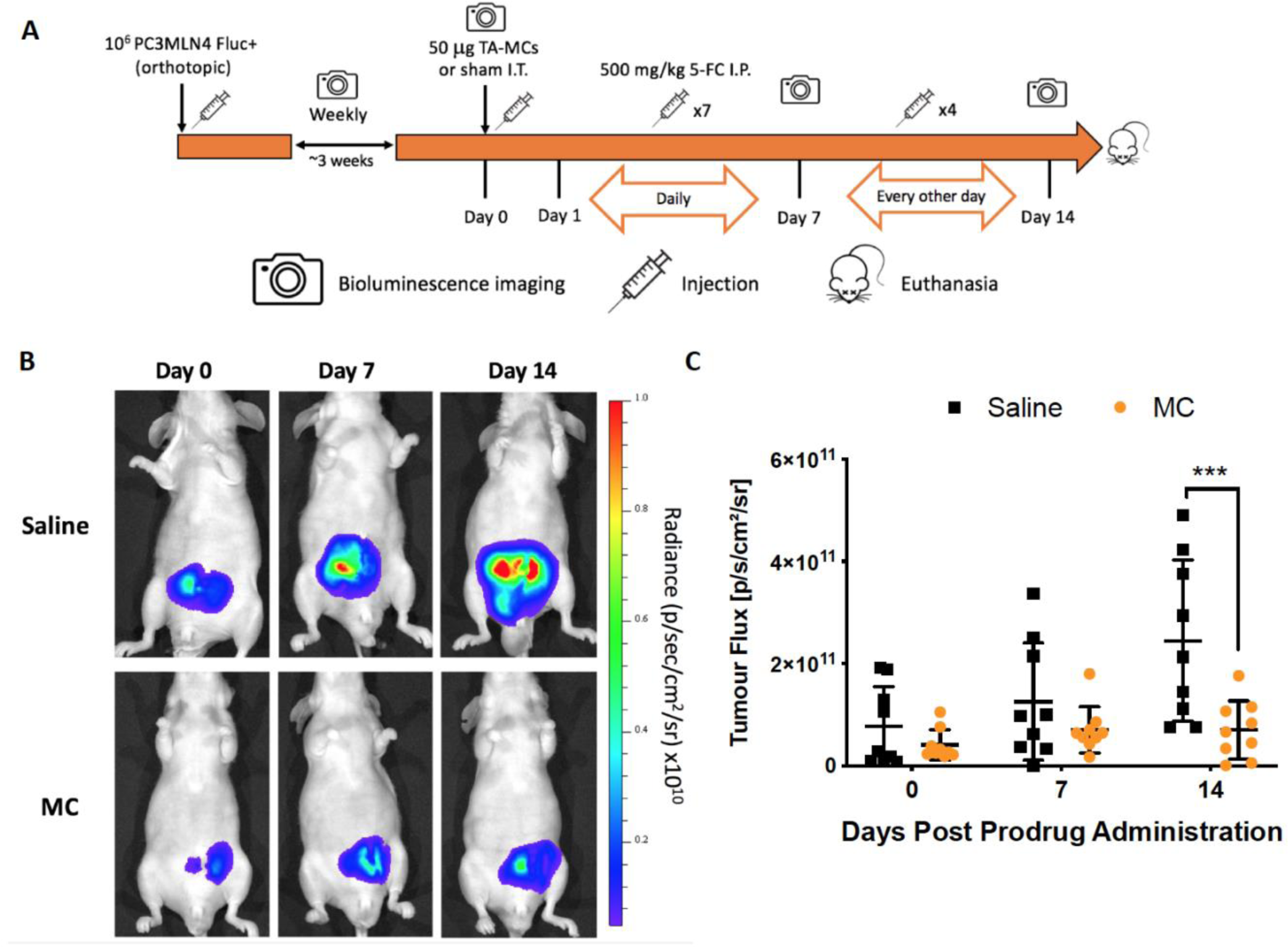
Assessment of therapeutic effect in nude mice with high-survivin orthotopic prostate tumours. (A) Timeline of intratumoural administration of pSurv-CD:UPRT-MCs, 5-FC treatment and BLI timepoints. (B) Representative FLuc BLI images with (D) quantification from nude mice with PC3MLN4 FLuc^+^ tumours post-injection of intratumoural saline or pSurv-CD:UPRT-MCs until endpoint at 14 days (n=9). Data are presented as mean ± SD (\*\*\**p* < 0.005).

## Discussion

Important characteristics for gene-based technologies for cancer detection and treatment are sufficient expression from tumours to produce a detectable signal or therapeutic effect, while also presenting minimal expression of transgenes in normal tissues. Survivin-driven, tumour-activatable MCs represent a promising platform for this purpose. Our initial work on this technology described activatable MCs encoding SEAP for detection of lung melanoma tumours [29]. This system was then adapted for PCa, where mice carrying high survivin, aggressive subcutaneous PCa tumours could be distinguished from low survivin, non-aggressive tumours via a blood SEAP reporter test [31]. Here, we report our latest iteration of the tumour-activatable MC technology, by introducing urinary GLuc reporter tests for tumour detection, and CD:UPRT-mediated GDEPT, describing our first combined tumour-activatable MC theranostic system.

We first sought to improve on blood-based diagnostic MCs by replacing SEAP with a bicistronic cassette encoding both GLuc and CD:UPRT. Despite initial assessments with these all-in-on MCs showing potential for simultaneous theranostics, we found greater efficacy when each transgene was encoded separately on individual MCs. GLuc-expressing MCs produced reporter levels titrated to survivin expression in transfected cells. Notably, these diagnostic MCs exhibited markedly higher output than their PP counterparts, likely attributed to both increased transfection efficiency and an improved expression profile [27], highlighting the value of MCs over plasmids in improving detection sensitivity. By measuring urine GLuc activity, diagnostic MCs injected intratumorally were able to specifically discern mice carrying aggressive, high-survivin PCMLN4 prostate tumours from mice with low-survivin LNCaP tumours, as well as healthy mice receiving intraprostatic MC injections. Due to the ease of sampling urine longitudinally, we were able to perform AUC analysis of urine GLuc activity measured daily over a week and found mice carrying aggressive tumours displayed significantly increased Gluc AUC compared to mice with LNCaP tumours and healthy mice. Estimating total GLuc output and assessing secretion kinetics over time may prove to be especially fruitful clinically, as these diagnostic metrics may be more accurate than a single, one-time measurement [42]. We also designed therapeutic MCs expressing CD:UPRT to be used in conjunction with our diagnostic MCs for treatment of GLuc-detected, survivin-high aggressive tumours. Therapeutic MCs selectively limited the *in vitro* growth of survivin-high PC3MLN4 cells but not survivin-poor LNCaP cells. Administered intratumourally, these therapeutic MCs attenuated the growth of aggressive prostate tumours over a 14-day period compared to sham-treated mice. By improving transfection efficiency and transgene expression, we hypothesize this therapeutic effect can be enhanced to reduce burden of aggressive tumours.

In this study, we chose PCa as our initial model to evaluate tumour-activatable MCs due to the unique diagnostic challenge that PCa presents, attributed to its high prevalence but varied lethality across individuals. PCa becomes lethal when it metastasizes outside the prostate, and we envision using diagnostic MCs to stratify patients with primary tumours into groups at high- and low-risk of future metastasis. This important prognostic information could improve patient-centred care and reduce overtreatment. Due to the transient nature of our episomal MCs, particularly in dividing cancer cells, GLuc will not be detectable in patients for extended periods of time, potentially allowing for repeated administration of GLuc-expressing MCs as a monitoring tool [43]. This synthetic reporter system may be used in conjunction with current diagnostic techniques like biopsy and prostate-specific antigen screening to improve disease management. Once patients with aggressive, high-risk PCa have been identified, therapeutic MCs may be used sequentially for tumour therapy and these MCs could even be delivered intraprostatically during a standard 12-core biopsy. Therapeutic MCs would act as an intermediate treatment post-diagnosis to prevent cancer spreading prior to patients receiving more radical procedures such as prostatectomy or brachytherapy, as these procedures are considerably more effective for early-stage disease [44]. A dual-MC theranostic system allows for separate administration of MCs pre- and post-diagnosis to match their clinical usage, whereas the same would not be possible for an all-in-one theranostic MC as both genes would be administered together. One potential alternative is to explore promoters such as astrocyte-elevated gene 1 (AEG-1) [45] or prostate specific membrane antigen (PSMA) [46], among others, which have been used for prostate cancer-specific transgene expression. Beyond prostate cancer, our MC system may be readily adapted as a pan-cancer theranostic technology as survivin is highly expressed in many common cancers including breast, lung, pancreatic, ovarian, colorectal, and liver [47].

Due to previous work showing that survivin expression correlates with increasing Gleason grade [16], pSurvivin was an attractive driver of transgene expression for our MCs. However, we also report some limitations of pSurvivin such as low levels of transgene expression in healthy tissues, and its weak activity compared to stronger, albeit constitutive, promoters. These drawbacks limit the sensitivity of our system, which could impact the viability of systemically delivered constructs in humans. One solution to increase transgene expression is to amplify pSurvivin activity using custom regulatory elements. For instance, the two-step transcriptional amplification system (TSTA) uses a GAL4-VP16 fusion protein and GAL4 DNA binding sites upstream of transgenes of interest to enhance tissue-specific expression from weaker promoters [48, 49] and these TSTA elements have yet to be included in MCs. The inclusion of a scaffold/matrix attachment region (S/MAR) into the 5’ untranslated region may also provide architectural anchors to promote replication of episomal gene vectors using host machinery [50]. S/MAR sequences have been shown to enhance and prolong MC expression in rapidly proliferating cells [51], and further *in vivo* study could establish S/MAR as a way to strengthen MC expression. Another important diagnostic use of tumour-activatable MCs is delivery of imaging reporter genes. Several groups have explored tumor-activatable vectors encoding imaging reporter genes for cancer detection with imaging modalities such as fluorescence [13], photoacoustic [52], and positron emission tomography (PET) [45]. Work using survivin-driven MCs to drive imaging reporter gene expression is ongoing in our group.

Many human clinical trials have utilized plasmid vectors to deliver transgenes [23, 53]. Beyond improved transfection rates compared to plasmids, MCs are more resistant to gene silencing due to fewer CpG motifs [54], DNA shearing forces [55], possessing a higher supercoiled fraction [56], and having enhanced serum stability [57]. To facilitate use in humans, a critical aspect to optimize is the MC to transfection agent ratio, and MCs allow for a higher effective dose than plasmids because a given transfection agent amount will contain more moles of MCs than plasmids. Another important factor to consider for translation is delivery. To maximize local delivery of MCs in this study, we performed intratumoural injections into multiple tumour loci. Despite some concerns such as leakage and uneven distribution, intraprostatic injections have already been performed for treatment of chronic prostatitis and benign prostatic hyperplasia [58] and for clinical trials with GDEPT for prostate cancer [59]. It would be valuable, nonetheless, to cultivate our system for systemic delivery. Our first work describing melanoma detection delivered SEAP-MCs intravenously and robustly identified mice bearing lung tumours by measuring blood SEAP activity [29]. The linear PEI agent we have used here primarily delivers non-viral vectors to the lungs and liver, and so systemic delivery to the prostate can be improved by using targeted transfection agents and/or nanoparticles [60, 61]. Exploring technologies to provide robust tumour-specific systemic delivery of MCs is a major focus going forward.

In this study, we built and evaluated a novel theranostic system comprised of a pair survivin-driven MCs for detection and treatment of aggressive prostate tumours. This system may serve as an effective way to identify prostate cancer patients with high-risk lesions and expedite treatment post-diagnosis, with the hope to ultimately lessen the psychophysical burden of this terrible disease.

## Abbreviations

AEG-1: astrocyte-elevated gene 1
BIRC5: baculoviral inhibitor of apoptosis repeat-containing 5
CD:UPRT: cytosine deaminase uracil phosphoribosyltransferase
DAB: 3,3-Diaminobenzidine
5-FC: 5-fluorocytosine
5-FUMP: 5-fluorouridine monphosphate
FLuc: *Firefly* luciferase
hNIS: human sodium iodine symporter
GDEPT: gene-directed enzyme prodrug therapy
GLuc: *Gaussia* luciferase
MC: minicircles
MR: magnetic resonance
OATP: organic anion transporting polypeptide
PCa: prostate cancer
PEI: polyethylenimine
PET: positron emission tomography
PP: parental plasmid
PSMA: prostate specific membrane antigen
RLuc: *Renilla* luciferase
SEAP: secreted embryonic alkaline phosphatase
S/MAR: scaffold/matrix attachment region
tdT: tdTomato
TSTA: two-step transcriptional amplification system.

## Acknowledgments

This work was in part funded by Canada Graduate Master’s Scholarship from the National Science and Engineering Research Council of Canada (TW). This work was also funded by a Movember Discovery Grant from Prostate Cancer Canada (JAR) and the Petro Canada Young Innovator Award from Western University (JAR).

## Contributions

This work was entirely supervised by JAR. Study was designed by TW and JAR. Data acquisition and analysis was performed by TW, DG, and YC. Results were interpreted by TW, YC, and JAR. Manuscript was prepared by TW, AA, and JAR. All authors have edited, reviewed, and approved of this submission.

