## Supplemental Figures for "A survivin-driven tumour-activatable minicircle system for prostate cancer theranostics"

**Supplementary Figures:**

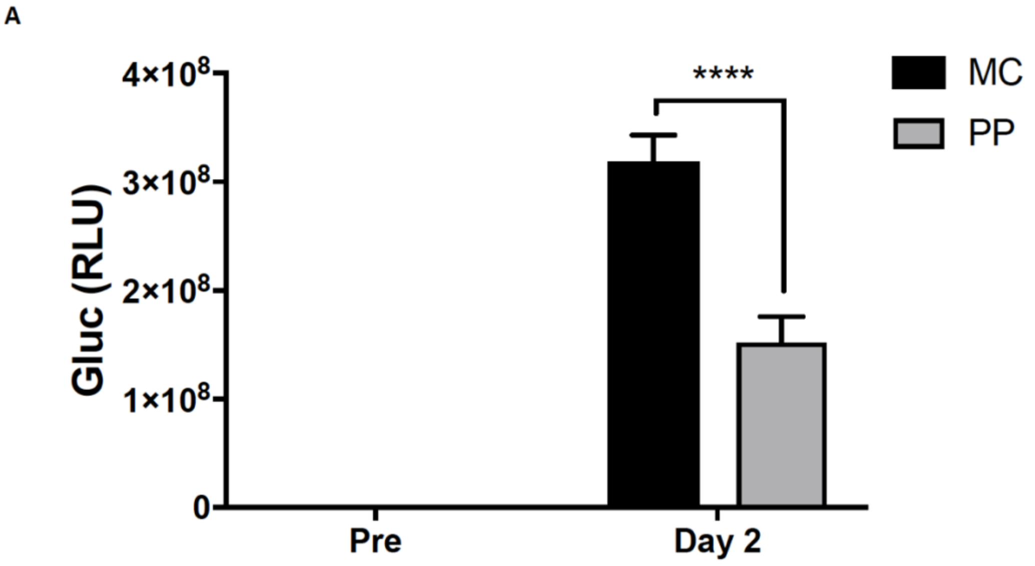


**Figure S1. Comparison of GLuc activity from PPs versus MCs.** GLuc activity in supernatant collected from PC3MLN4 cells transfected with equimolar pSurv-GLuc-PP or MC pre- and on day 2 post-transfection (n=5). Data are presented as mean ± SD (*****p* < 0.001).

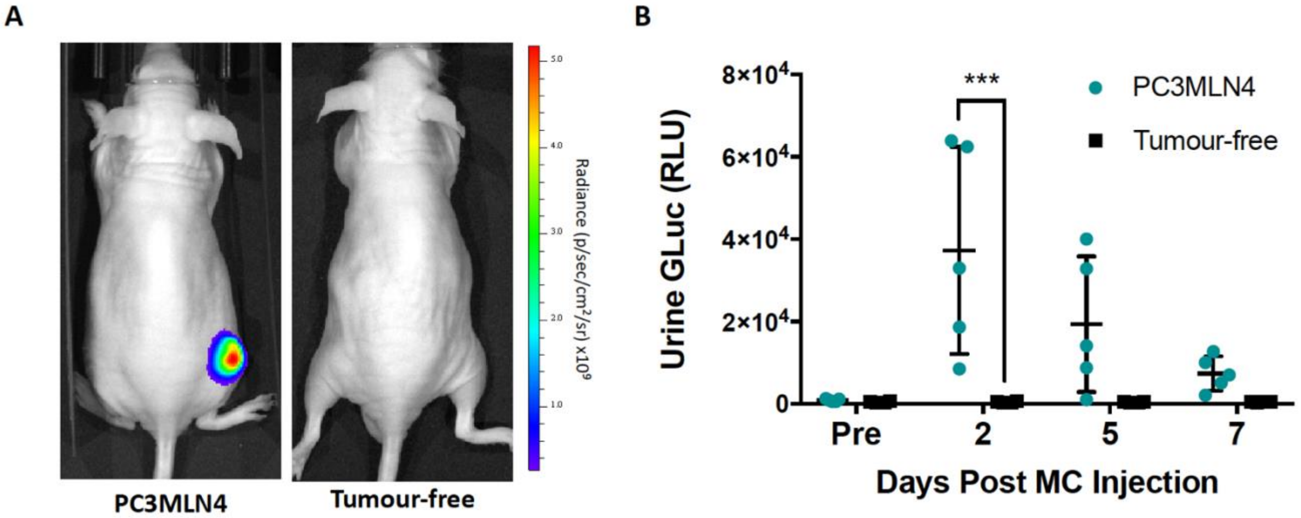


**Figure S2. Evaluation of urine GLuc activity in nude mice with subcutaneous prostate cancer tumours.** (A) Representative FLuc BLI images of mice with subcutaneous tumours 30 days after tumour implantation and tumour-free animals. (B) Urine GLuc activity post intratumoural MC administration from mice bearing PC3MLN4 tumours (n=5) or post intramuscular injection from tumour-free mice (n=4). Data are presented as mean ± SD (****p* < 0.005).

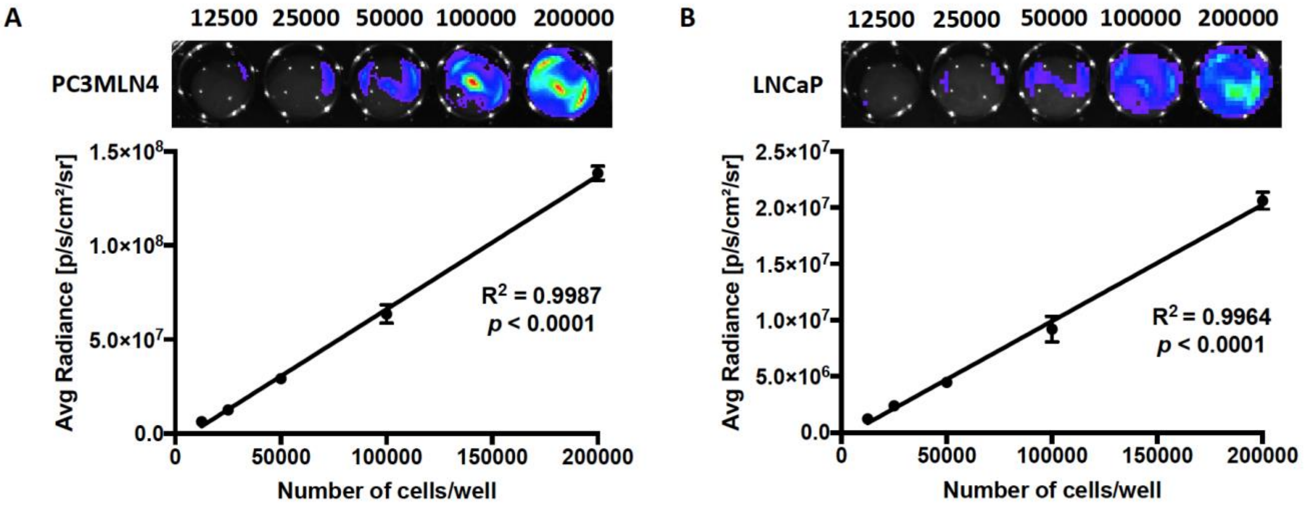


**Figure S3. Assessment of cellular FLuc signal as indicator of cell viability.** Bioluminescence imaging of (A) PC3MLN4 FLuc+ and (B) LNCaP FLuc+ cells seeded at increasing densities (n=3). Images were taken one day after seeding. Data are presented as mean ± SD.

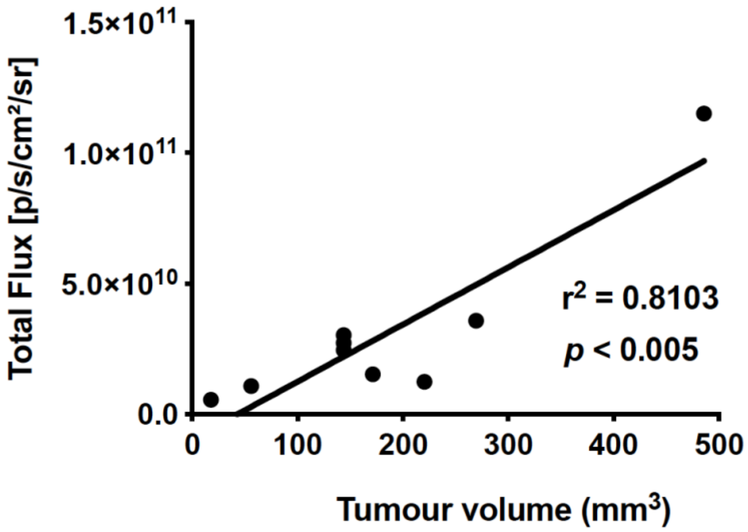


**Figure S4. Assessment of tumour volume versus BLI signal.** BLI signal of orthotopic PC3MLN4 FLuc+ tumours taken one day prior to measurement of tumour dimensions using a manual caliper (n=9).

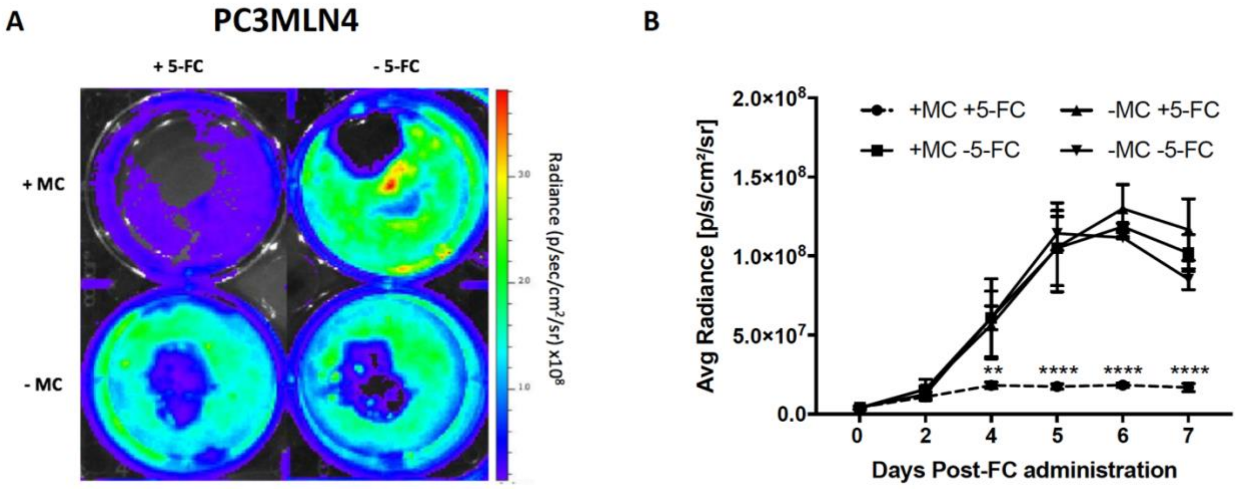


**Figure S5. Characterization of suicide gene therapy system on FLuc-expressing PCa cells.** (A) Bioluminescence images of PC3MLN4 FLuc+ cells on day 6 post-transfection with pSurv-CD:UPRT-MCs with (B) quantification (n=3). Data are presented as mean ± SD.
